# RNA-binding activity of TRIM25 is mediated by its PRY/SPRY domain and is required for ubiquitination

**DOI:** 10.1101/200410

**Authors:** Nila Roy Choudhury, Gregory Heikel, Maryia Trubitsyna, Peter Kubik, Jakub Stanislaw Nowak, Shaun Webb, Sander Granneman, Christos Spanos, Juri Rappsilber, Alfredo Castello, Gracjan Michlewski

## Abstract

TRIM25 is a novel RNA-binding protein and a member of the Tripartite Motif (TRIM) family of E3 ubiquitin ligases, which plays a pivotal role in the innate immune response. Almost nothing is known about its RNA-related roles in cell biology. Furthermore, its RNA-binding domain has not been characterized. Here, we reveal that RNA-binding activity of TRIM25 is mediated by its PRY/SPRY domain, which we postulate to be a novel RNA-binding domain. Using CLIP-seq and SILAC-based co-immunoprecipitation assays, we uncover TRIM25’s endogenous RNA targets and protein binding partners. Finally, we show that the RNA-binding activity of TRIM25 is important for its ubiquitin ligase function. These results reveal new insights into the molecular roles and characteristics of RNA-binding E3 ubiquitin ligases and demonstrate that RNA could be an essential factor for their biological functions.

## INTRODUCTION

More than 300 novel RNA-binding proteins have been identified in HeLa (Castello et al., 2012), HEK293 (Baltz et al., 2012) and mouse ES cells (Kwon et al., 2013). Furthermore, a recent study identified 393 new RNA-binding proteins unique to cardiomyocytes (Liao et al., 2016). Strikingly, many of these proteins were not previously known for their RNA-binding properties and do not contain canonical RNA-binding domains. Intriguingly, E3 ubiquitin ligases from the Tripartite Motif (TRIM) family (TRIM25, TRIM56 and TRIM71) are among these newly identified RNA-binding proteins. TRIM proteins have a characteristic architecture that includes RING zinc-finger, B-Box, coiled-coil (CC) and variable C-terminal domains. There are over 80 TRIM proteins in the human genome, and they have been divided into 11 subfamilies based on their structural composition (Versteeg et al., 2013). The largest subfamily (C-IV) of 33 TRIM proteins is characterized by the PRY/SPRY domain at the C-terminal end. Most the proteins from the TRIM family are involved in innate immune responses (Jefferies et al., 2011; Rajsbaum et al., 2014; van Tol et al., 2017). For example, TRIM25 plays a role in the RIG-I-mediated interferon response to viral RNAs (Ozato et al., 2008; Versteeg et al., 2013). RIG-I recognizes viral RNA molecules with a 5’-triphosphate (5’-ppp-RNA) (Yoneyama et al., 2004) and initiates a downstream signaling cascade that culminates with the expression of type I interferons (interferon α and β) (Loo and Gale, 2011) and an anti-viral response. TRIM25 mediates K63-linked ubiquitination of RIG-I, which leads to efficient recruitment of its downstream partners. Such recruitment in turn triggers the interferon response (Gack et al., 2007). Recently, TRIM25 was shown to be required for the antiviral activity of RNA-binding Zinc Finger Antiviral Protein (ZAP) (Li et al., 2017; Zheng et al., 2017). Almost nothing is known about the RNA-mediated roles and characteristics of these RNA-binding E3 ligases.

Although TRIM25 has been generally well characterized, the functional importance of its RNA-binding activity remains unclear. Our recent study provided first clues: we have shown that the binding of TRIM25 to pre-let-7 leads to its more efficient Lin28a-mediated uridylation (Choudhury et al., 2014). This finding raised the possibility that RNA processing may be spatially controlled through post-translational protein modifications by RNA-binding protein-modifying enzymes, such as TRIM25. Furthermore, TRIM25 could be using RNA as a scaffold to ubiquitinate its targets. Importantly, the transcriptome and proteome-wide targets of TRIM25 have not yet been identified. Finally, its RNA-binding domain has not been unequivocally described.

Here, using data from a proteome-wide analysis of RNA-binding domains, RNA pull-downs and EMSA, we show that the PRY/SPRY domain of TRIM25 harbors its RNA-binding activity. Furthermore, TRIM25 CLIP-seq and quantitative SILAC-based co-immunoprecipitation assays reveal a comprehensive overview of interacting RNAs and proteins, respectively. Our CLIP-seq data identifies hundreds of coding and non-coding RNAs bound by TRIM25. We show that TRIM25 binds a distinct set of RNAs and prefers to bind sequences rich in G and C. We also demonstrate that TRIM25 associates with many RNA-binding proteins, thus suggesting it could play a role in regulating RNA metabolism. Crucially, we present evidence that the binding of TRIM25 to RNA is important for its ubiquitin ligase activity towards itself (autoubiquitination) and physiologically relevant target ZAP. Finally, by showing that the PRY/SPRY regions of other TRIM proteins rescue TRIM25’s RNA-binding activity, we propose that TRIM25 represents a novel group of E3 ligases with RNA-binding activity that use PRY/SPRY as a novel RNA-binding domain. These results provide original insights into the RNA-mediated activities of RNA-binding E3 ubiquitin ligases.

## RESULTS

### TRIM25 directly interacts with RNA through the PRY/SPRY domain

TRIM25 is known to associate with RNAs in cells and in cell extracts (Choudhury et al., 2014; Kwon et al., 2013). However, whether this interaction is direct or indirect remains unknown. To test this, we performed an electrophoretic mobility shift assay (EMSA) with pre-let-7a-1 and purified recombinant human His-tagged TRIM25, expressed in *E. coli* (Figure S1a). EMSA, with increasing amounts of His-TRIM25, revealed that TRIM25 interacts with RNA directly (Fig. 1a), with an observed Kd of 800 nM. This showed that TRIM25 was a *bona fide* RNA-binding protein.

**Fig. 1.**
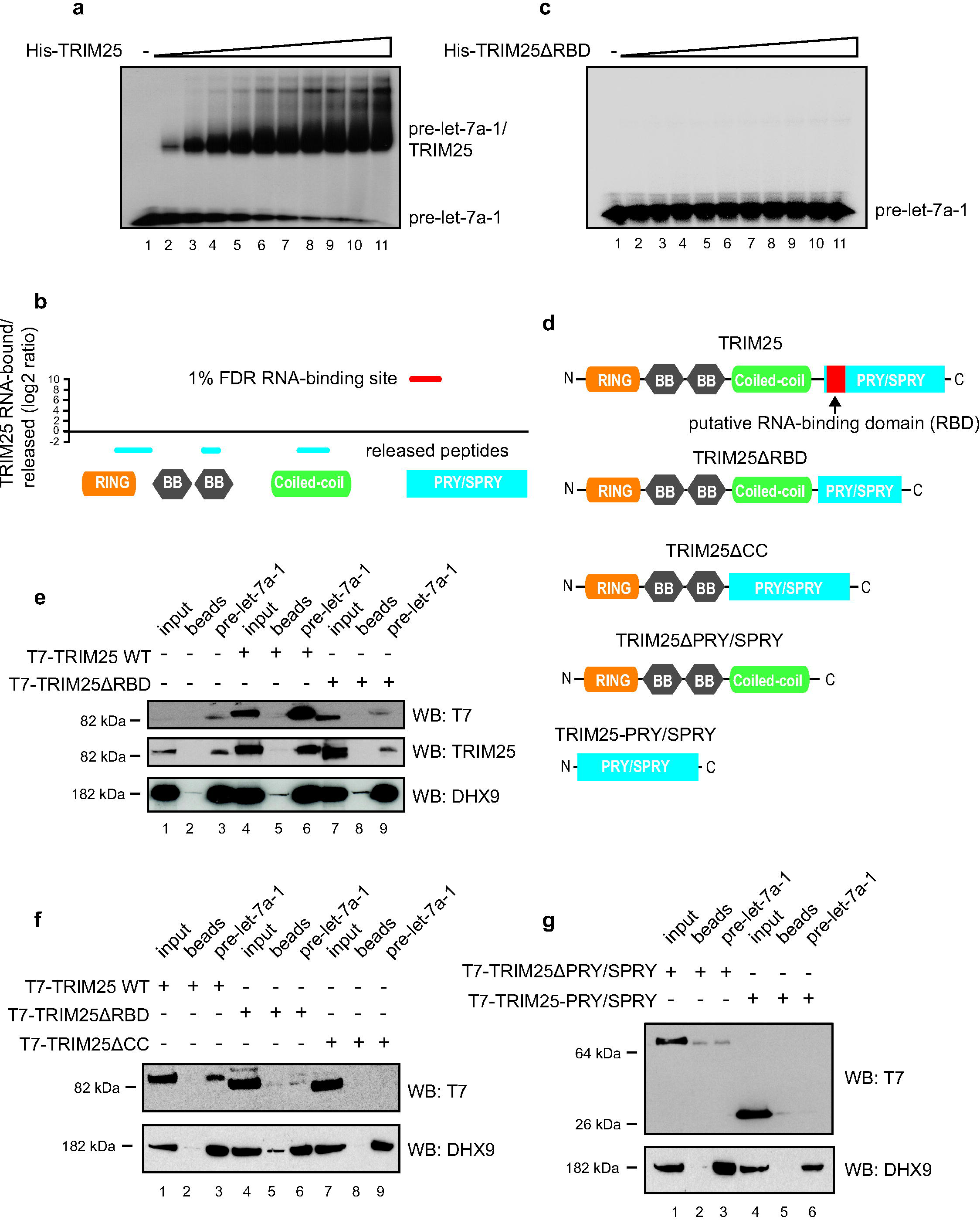
TRIM25 is a *bona fide* RNA-binding protein and the PRY/SPRY domain is responsible for binding to RNA. **(a)** EMSA analysis of recombinant His-tagged TRIM25 with pre-let-7a-1. Lane 1 represents the loading control. Lanes 2 to 11 show EMSA with increasing amounts of TRIM25 (200 ng, 400 ng, 600 ng, 800 ng, 1000 ng, 1200ng, 1400ng, 1600 ng, 1800 ng and 2000ng). **(b)** RNA capture assay result for TRIM25. Red line represents peptide enriched in RNA-bound fraction. Blue lines show peptides depleted from the RNA-bound fraction. **(c)** EMSA analysis of recombinant His-tagged TRIM25ΔRBD with pre-let-7a-1. Lane 1 represents the loading control. Lanes 2 to 11 show EMSA with increasing amount of TRIM25ΔRBD (200 ng, 400 ng, 600 ng, 800 ng, 1000 ng, 1200ng, 1400ng, 1600 ng, 1800 ng and 2000ng). **(d)** Domain architecture of wild type TRIM25 and deletions mutants. Relative position of RNA-enriched peptide is shown in red. **(e)** Western blot analyses against T7, TRIM25 and DHX9 of pre-let-7a-1 pull downs with HeLa cell extracts overexpressing T7-TRIM25 or T7-TRIM25ΔRBD. Lanes 1, 4 and 7 represent 4% (40 μg) of the loading controls (input). Lanes 2, 5 and 8 represent beads only pull downs. Lanes 3, 6 and 9 show pre-let-7a-1 pull down. Note that T7 antibody is highlighting an unspecific band visible in lanes 3 and 8. **(f)** Western blot analyses against T7 and DHX9 of pre-let-7a-1 pull downs with HeLa TRIM25 KO cell extracts overexpressing T7-TRIM25, T7-TRIM25ΔRBD or T7-TRIM25ΔCC. Lanes 1, 4 and 7 represent 4% (40 μg) of the loading controls (input). Lanes 2, 5 and 8 represent beads only pull downs. Lanes 3, 6 and 9 show pre-let-7a-1 pull down. **(g)** Western blot analyses against T7 and DHX9 of pre-let-7a-1 pull downs with HeLa TRIM25 KO cell extracts overexpressing T7-TRIM25ΔPRY/SPRY or T7-TRIM25-PRY/SPR. Lanes 1 and 4 represent 4% (40 μg) of the loading controls (input). Lanes 2 and 5 represent beads only controls. Lanes 3 and 6 show pre-let-7a-1 pull down.

Previously, TRIM25 has been shown to bind RNA through its coiled-coil CC domain (Kwon et al., 2013), which is responsible for its dimerization and oligomerization (Koliopoulos et al., 2016; Li et al., 2014; Sanchez et al., 2014; Streich et al., 2013). Surprisingly, a proteome-wide analysis of RNA-binding domains (RBDs) in HeLa cells revealed that the TRIM25 PRY/SPRY domain may be responsible for direct binding to RNA (Fig. 1b) (Castello et al., 2016). In brief, this method combines UV protein-RNA crosslinking, oligo(dT) capture and controlled proteolysis for a high-resolution (∼17 amino acids on average) delineation of the protein region engaged in RNA binding. In the case of TRIM25, the peptide enriched in the RNA-bound fraction mapped to the PRY/SPRY domain (TRIM25 residues 470-508), demonstrating direct RNA binding (red line; 1% false discovery rate (FDR), Fig. 1b). In contrast, peptides mapping to other regions of the protein, including the CC domain, were released into the supernatant after the protease cleavage (blue lines), suggesting a lack of RNA-binding activity (Fig. 1b). To validate this, we expressed and purified a recombinant His-TRIM25 deletion mutant (lacking residues 470-508 located in the PRY/SPRY domain) (Figure S1a), which we named TRIM25ΔRBD (RBD - RNA Binding Domain), and tested its binding activity *in vitro* (Fig. 1c-1d). EMSA analysis with increasing amounts of His-TRIM25ΔRBD revealed no shift in the pre-let-7a-1 substrate (Fig. 1c). Likewise, RNA pull-down assays in HeLa cell extracts showed that endogenous and T7-tagged TRIM25 were efficiently pulled down with pre-let-7a-1, whereas T7-tagged TRIM25ΔRBD was not (Fig. 1e). These results strongly indicate that the TRIM25 PRY/SPRY domain harbors its RNA-binding activity.

Importantly, some RNA-binding proteins must dimerize to be able to bind RNA (Feracci et al., 2016; Lunde et al., 2007). Since TRIM25 is known to dimerize and tetramerize through its CC domain with minimal contributions from the PRY/SPRY (Koliopoulos et al., 2016; Li et al., 2014; Sanchez et al., 2014; Streich et al., 2013), we decided to generate TRIM25 knockout (KO) cells, which removed any confounding effects of endogenous TRIM25. To obtain TRIM25 KO cells we used CRISPR/Cas9 targeting followed by clonal selection and dot blot analysis. This strategy yielded several HeLa cell clones with complete loss of TRIM25 expression (Figure S2a). For further analysis, we selected clone C3 (Figure S2b). Next, we performed RNA pull-down assays in extracts from the TRIM25 KO cells with T7-TRIM25ΔRBD and additional TRIM25 truncations (Fig. 1d). These experiments confirmed that neither T7-TRIM25ΔRBD nor T7-TRIM25ΔCC retained RNA-binding activity (Fig. 1f). Importantly, neither T7-TRIM25ΔPRY/SPRY nor T7-TRIM25-PRY/SPRY could bind RNA (Fig. 1g). This shows that the TRIM25 CC domain is essential, whereas the isolated PRY/SPRY domain is not sufficient for binding to RNA. These results suggest that TRIM25 binding to RNA could be dependent on its CC-mediated dimerization and are in line with previous results, which linked the CC domain with TRIM25’s RNA-binding activity (Kwon et al., 2013).

Finally, we wanted to check if TRIM25ΔRBD could fold correctly and form multimeric complexes. To test this, we overexpressed T7-tagged TRIM25 with EGFP-tagged TRIM25 (with or without the ΔRBD mutation) in TRIM25 KO cells and performed anti-T7 co-immunoprecipitation (co-IP) assays (Fig. 2a). Our results showed that both wild-type T7-TRIM25 and T7-TRIM25ΔRBD could interact with EGFP-TRIM25 (Fig. 2a). The control experiment revealed that the T7 antibody co-IPed T7-tagged but not EGFP-tagged TRIM25 (Fig. 2b). Moreover, we confirmed that T7-TRIM25ΔCC or the isolated T7-TRIM25-PRY/SPRY domain could not form multimeric complexes with EGFP-TRIM25 (Fig. 2c). Importantly, both the wild-type and TRIM25ΔRBD recombinant proteins displayed a similar Tm (43 C^o^) in a thermal denaturation assay (Figure S1b). Furthermore, using Size Exclusion Chromatography with Multi-Angle Light Scattering (SEC-MALS), we established that there was no difference in the elution profiles between the wild-type and the mutant protein (Figure S1c). The MALS-measured mass distribution for His-TRIM25 was 263 ± 5.3 kDa, with an Mw/Mn of ∼ 1.01. The His-TRIM25ΔRBD had values of 258 ± 5 kDa, and ≥ 1.003 Mw/Mn. This suggests that both wild type and TRIM25ΔRBD formed tetramers in our preparation. Taken together, these results show that TRIM25 harboring a ΔRBD mutation still supports interactions *in trans* with another TRIM25 molecule and that the ΔRBD mutation does not influence the overall folding or multimerization of the TRIM25 protein.

**Fig. 2.**
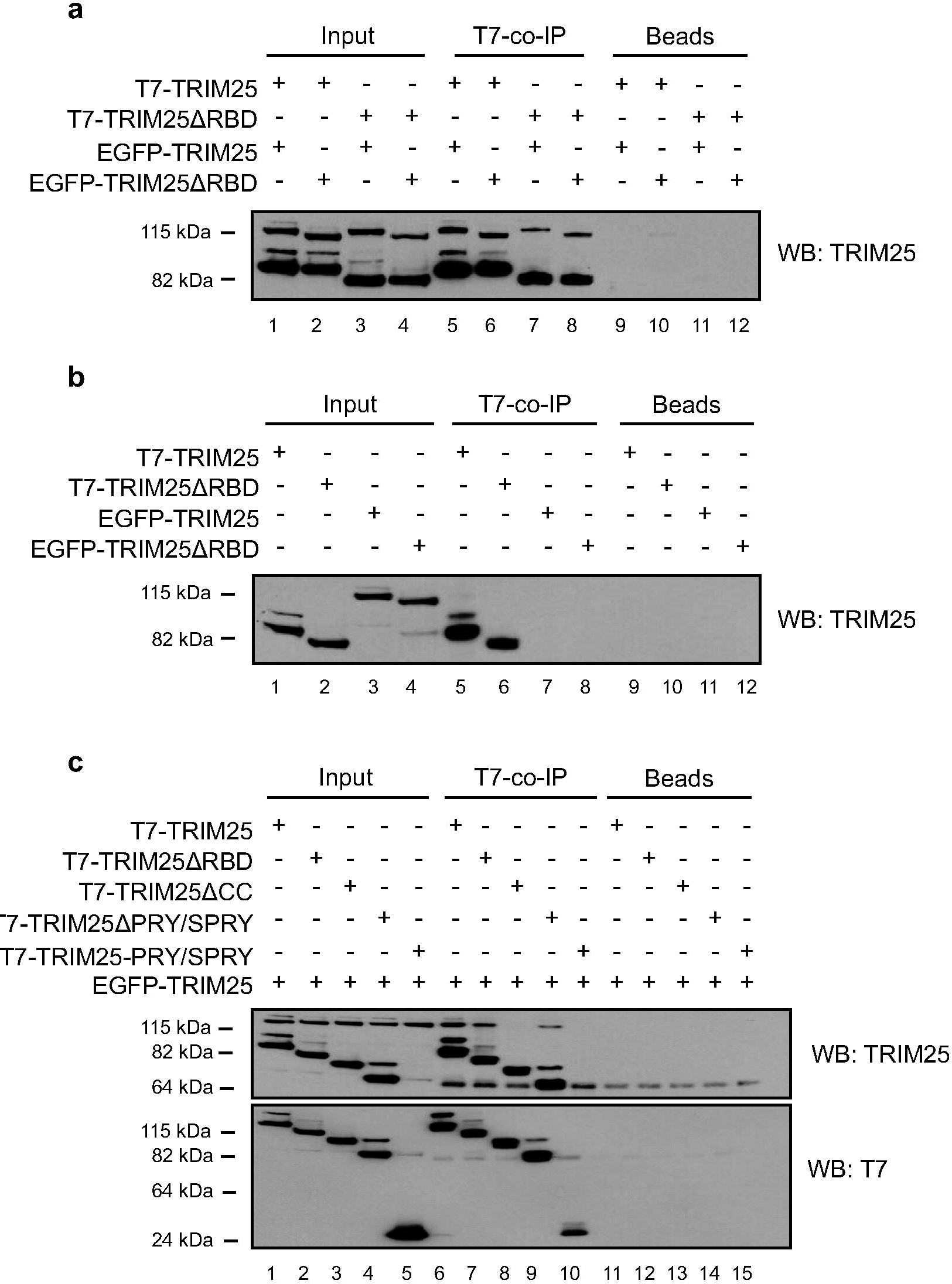
T7-TRIM25ΔRBD supports TRIM25 dimerization. **(a)** Co-IP between T7-tagged TRIM25 and EGFP-tagged TRIM25 in cell extracts prepared from HeLa TRIM25 KO cells overexpressing wild type or ΔRBD TRIM25 mutants. Lanes 1-4 represent loading controls. Lanes 5-8 show co-immunoprecipitations with anti-T7 antibody. Lanes 9-12 represent control co-immunoprecipitations with IgG antibody bound to protein-A beads. The bound proteins were analyzed by western blotting with anti-TRIM25 antibody. **(b)** Control co-IP shows that EGFP-TRIM25 and EGFP-TRIM25 are not co-IPed by anti-T7 antibody. Lanes 1-4 represent loading controls. Lanes 5-8 show co-immunoprecipitations with anti-T7 antibody. Lanes 9-12 represent control co-immunoprecipitations with IgG antibody bound to protein-A beads. The bound proteins were analyzed by western blotting with anti-TRIM25 antibody. **(c)** Control co-IP in HeLa TRIM25 KO cell extracts reveals that T7-TRIM25ΔRBD and T7-TRIM25ΔPRY/SPRY but not T7-TRIM25ΔCC or T7-TRIM25-PRY/SPRY support dimerization. Lanes 1-4 represent loading controls. Lanes 5-8 show co-immunoprecipitations with anti-T7 antibody. Lanes 9-12 represent control co-immunoprecipitations with IgG antibody bound to protein-A beads. The bound proteins were analyzed by western blotting with anti-TRIM25 and anti-T7 antibodies.

### TRIM25 co-immunoprecipitates with proteins involved in RNA metabolism

To find TRIM25-interacting proteins, we performed co-IPs of T7-tagged TRIM25 in HeLa cells coupled with quantitative SILAC mass spectrometry. In brief, wild-type T7-tagged TRIM25 was expressed in HeLa cells, and the co-IP was performed using protein A agarose beads coupled with the anti-T7 antibody (Fig. 3a). The control cells (without T7-TRIM25) were grown in “heavy” medium, and the cells expressing T7-tagged TRIM25 were grown in “light” SILAC medium. This set-up allowed for a quantitative assessment of proteins that bound specifically to TRIM25. After the mass spectrometry analysis, the most enriched protein in the co-IP of T7-TRIM25-expressing cells was TRIM25, which validated our approach (Table S1). Next, we performed functional GO-term annotations for enriched proteins using DAVID (Huang da et al., 2009) and visualized the results in Cytoscape with the Enrichment Map Plugin (Merico et al., 2010). Importantly, we found many proteins involved in RNA metabolism, such as helicases, RNA processing, nucleotide binding, ribosomal proteins or proteins involved in RNA stability (Fig. 3a). Recently, a large-scale protein-protein interactome analysis revealed that RNA-binding proteins are likely to co-immunoprecipitate higher numbers of other RNA-binding proteins (Brannan et al., 2016). Thus, our results emphasize the role of TRIM25 as an RNA-binding protein.

**Fig. 3.**
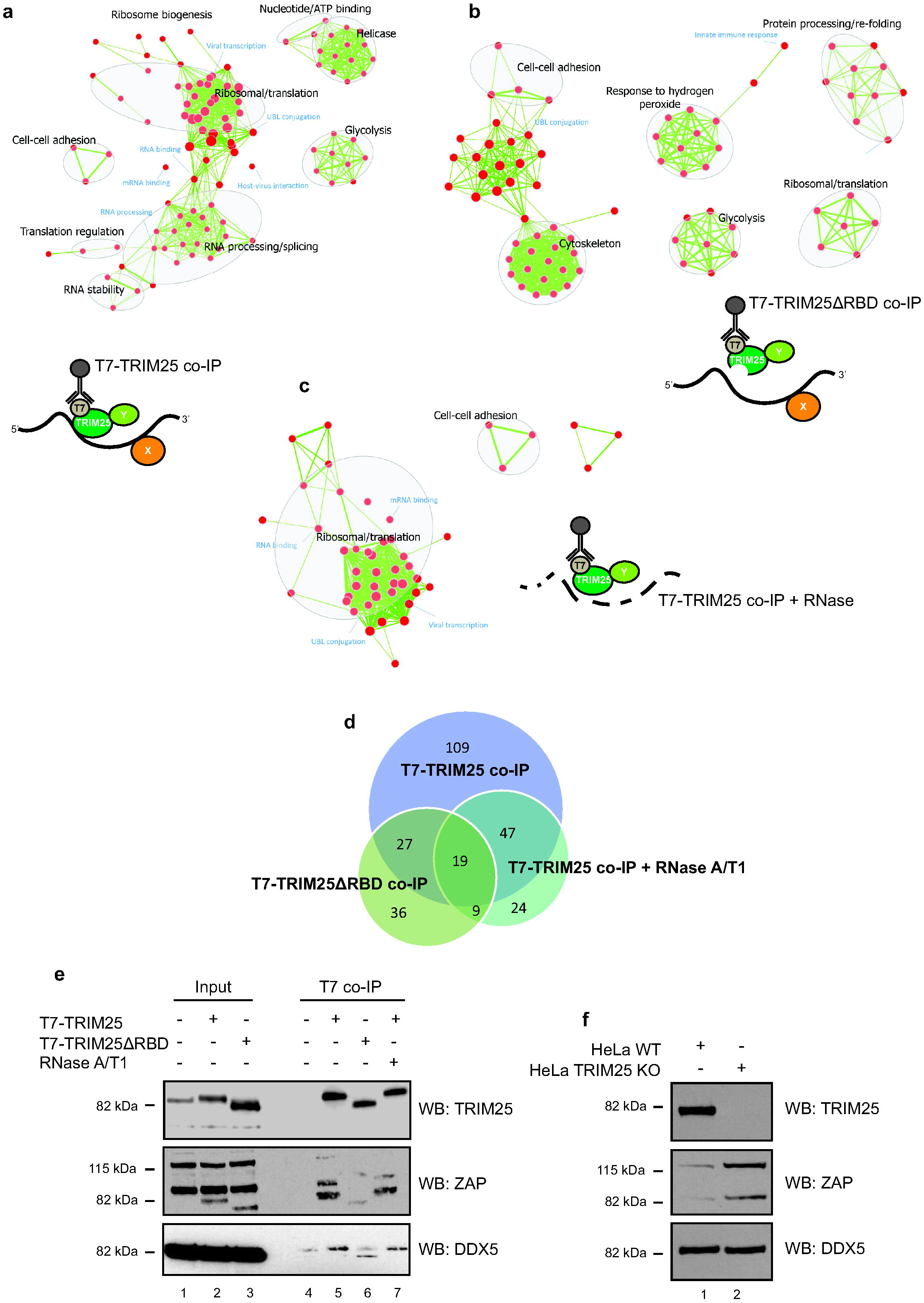
SILAC combined with TRIM25 co-immunoprecipitation reveals a network of associated RNA-binding proteins. **(a)** A diagram of a co-IP experiment with T7-tagged TRIM25 in the presence of endogenous RNAs together with Gene Ontology enrichment map of proteins co-immunoprecipitated with T7-TRIM25. Enriched protein sets are represented as nodes (red circles) connected by edges (green lines). With the size of nodes and edges representing the number of proteins in gene sets and amount of overlap. **(b)** A diagram of a co-IP experiment with T7-tagged TRIM25ΔRBD in the presence of endogenous RNAs accompanied by the Gene Ontology enrichment map of proteins co-immunoprecipitated with T7-TRIM25ΔRBD. **(c)** A diagram of a co-IP experiment with T7-tagged TRIM25 in the presence RNases together with Gene Ontology enrichment map of proteins co-immunoprecipitated with T7-TRIM25 in the presence of RNases. **(d)** Venn diagram representing numbers of overlapping proteins identified in the co-immunoprecipitations with T7-TRIM25, T7-TRIM25ΔRBD and T7-TRIM25 in the presence of RNases. **(e)** Validation of co-IP experiments show western blot against ZAP and DDX5 proteins. ZAP has two isoforms which are co-IP by T7-TRIM25 but not by T7-TRIM25ΔRBD or T7-TRIM25 in the presence of RNases. Note that anti-ZAP antibody is picking up signal from overexpressed TRIM25. DDX5 binding to TRIM25 is partially RNA-dependent as co-IP by T7-TRIM25 in the presence of RNases decreases but not eliminates its interaction with TRIM25. Lanes 1, 2 and 3 represent loading controls. Lane 4 shows control co-immunoprecipitations in mock HeLa cell extracts. Lanes 5,6 and 7 show anti-T7 co-IP. (**f**) Western blot analysis of wild type (WT) and TRIM25 KO HeLa cells shows upregulation of ZAP protein in the KO cells. DDX5 levels do not change and serve as a loading control.

Next, to see which of these protein-protein interactions are mediated by RNA, we performed the same co-IP assays using T7-tagged TRIM25ΔRBD (Fig. 3b) or T7-tagged TRIM25 but in the presence of RNases (Fig. 3c). In total, 109 proteins were absent in either the T7- TRIM25ΔRBD or RNase-treated T7-TRIM25 co-IPs when compared with the wild-type T7-TRIM25 co-IP (Fig. 3d). These proteins, which most likely interact with TRIM25 through RNA, are associated with RNA processing and splicing, RNA stability, RNA transport and helicase activity. We have validated that a recently identified, physiologically relevant TRIM25 ubiquitination substrate – RNA-binding Zinc-Finger Antiviral Protein (ZAP) (Li et al., 2017; Zheng et al., 2017) – interacts with T7-TRIM25 but not T7-TRIM25ΔRBD (Fig. 3e). ZAP has longer and shorter isoforms (ZAPL – long, ZAPS – short), which differ by presence or absence of poly(ADP-ribose) polymerase domain in the C-terminal region. The binding of both isoforms to TRIM25 was severely reduced upon RNase treatment proving that the interaction is dependent on the RNA (Fig. 3e). Likewise, DDX5 was predominantly detected in the co-IP with T7-TRIM25 but not T7-TRIM25ΔRBD, although the RNase treatment resulted in only a decreased interaction (Fig. 3e). Importantly, many proteins that were enriched in the T7-TRIM25ΔRBD and RNase-treated T7-TRIM25 assays were associated with ribosomal and translation functions (Fig. 3b-3c). They represent RNA-independent interactions with TRIM25 (Table S1). Surprisingly, the T7-TRIM25ΔRBD and RNase-treated T7-TRIM25 co-IPs detected several proteins that were not seen in the wild-type T7-TRIM25 co-IP. This could be due to a synthetic effect of the mutation, lack of binding to RNA or unspecific binding events. Altogether, these results reaffirm that the PRY/SPRY domain is important for RNA-binding and show that TRIM25 interacts with many RNA-binding proteins, using RNA as a scaffold.

We set out to find out if TRIM25 KO results in global perturbation of protein levels. To do so we used SILAC mass spectrometry on the wild type and TRIM25 KO HeLa cells. Our results showed that the levels of only a few proteins were affected more than twofold (Table S2). Among the affected proteins we found ZAP, which was substantially upregulated in TRIM25 KO cells when compared with the wild type cells. Western blot analysis of the wild type and KO cells validated this observation (Fig. 3f). To see if TRIM25-dependent proteasome degradation of ZAP contributes to its lower levels in the wild type cells we treated the cells with proteasome inhibitor MG132. Upon MG132 treatment we noticed increase of ZAP levels only in the wild type cells but not in the TRIM25 KO cells (Figure S2c). The signal corresponding to ubiquitin was elevated in both wild type and TRIM25 KO cells (Figure S2d). Importantly, overexpression of T7-TRIM25 but not T7-TRIM25ΔRBD or catalytically inactive T7-TRIM25ΔRING decreased the levels of ZAP in TRIM25 KO cells (Figure S2e). These results demonstrate that TRIM25-mediated proteasome degradation of ZAP contributes to its normal levels. Furthermore, these results suggest that the functional interaction between TRIM25 and ZAP is RNA-dependent.

### CLIP-seq reveals an extensive TRIM25 RNA interactome

To detect endogenous RNA substrates of TRIM25, we performed UV crosslinking and immunoprecipitations in HeLa cells transiently expressing T7-tagged TRIM25, followed by high-throughput sequencing of TRIM25-associated RNAs (CLIP-seq). After UV crosslinking and immunoprecipitation, we observed a specific RNA-protein complex (Figure S3a). CLIP-seq of T7-TRIM25ΔRBD resulted in a lack of detectable RNA-protein interactions (data not shown). The RNA from the T7-TRIM25/RNA complex was isolated using a CLIP-seq protocol and sequenced on a HiSeq platform. Data analysis and cluster identification using pyCRAC (Webb et al., 2014) revealed a significant correlation between three biological replicates with a correlation coefficient between clusters from 0.94 to 0.96 (Figure S4). Five thousand five hundred and forty-nine clusters were common between all three experiments, and we identified 2611 distinct transcripts (Table S3). Importantly, there was no correlation between transcript abundance, measured by a separate RNA-seq assay, and the cluster intensities, with a Pearson correlation of 0.1 (Figure S5). This shows that TRIM25 does not simply bind the most abundant RNAs but has a substrate preference. Among the identified transcripts, mRNAs and lincRNAs were the most abundant (Fig. 4a). GO-term annotation of TRIM25-bound transcripts revealed several nodes including RNA processing and translation, WD repeats, phosphorylation and ubiquitination (Figure S6a).

**Fig. 4.**
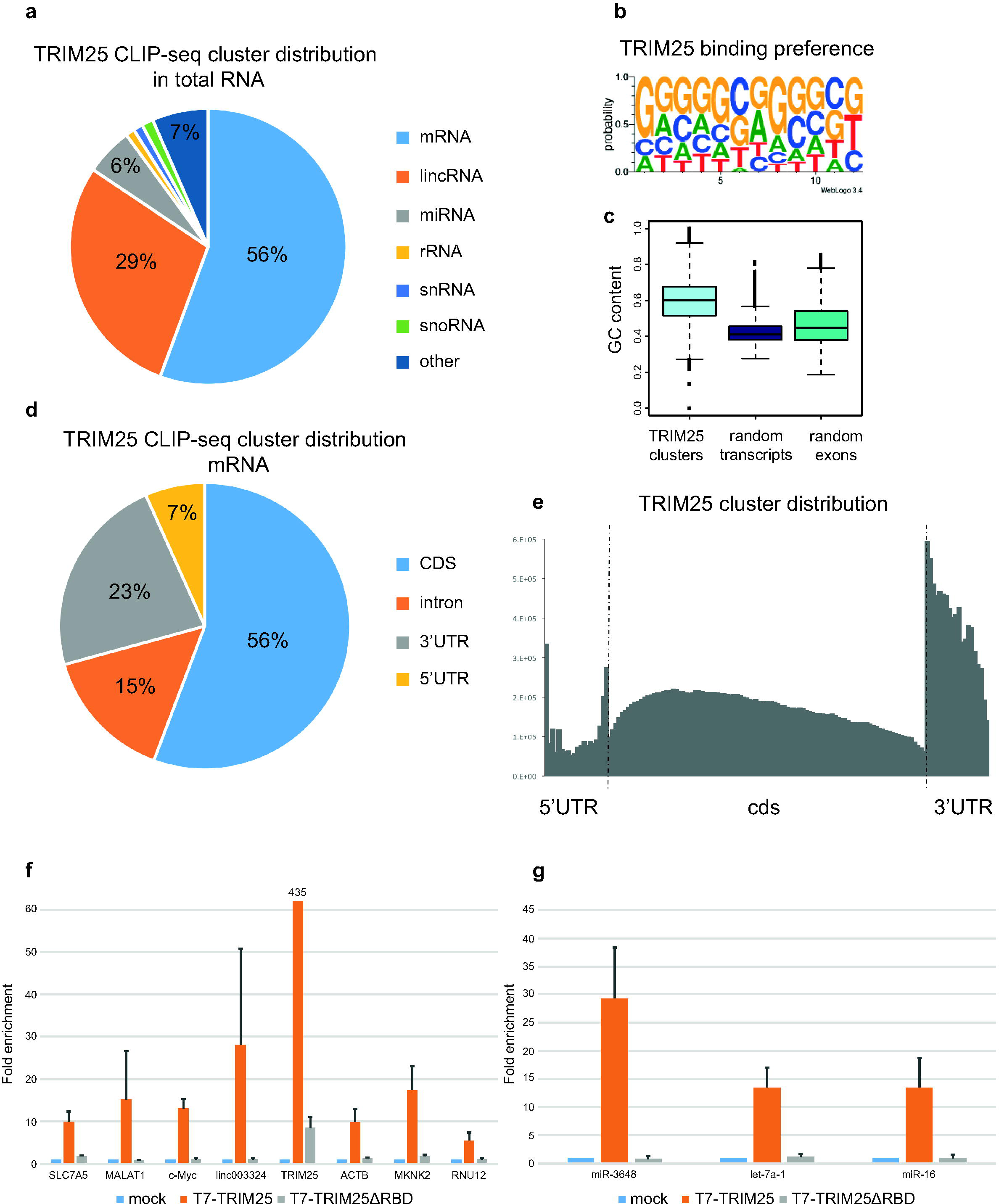
CLIP-seq of T7-TRIM25 in HeLa cells reveals novel *TRIM25* mRNA targets. **(a)** The distribution of TRIM25 CLIP-seq clusters in total RNA **(b)** The ab initio-derived TRIM25-binding sequence preference. T is representative of U in the RNA. **(c)** GC content is significantly enriched in TRIM25 CLIP-seq clusters as compared to randomly selected transcripts or exons with similar length distribution (transcripts detected in HeLa RNA-seq) -p-value 2.2e-16. **(d)** The distribution of TRIM25 CLIP-seq clusters in mRNAs. **(e)** The distribution of TRIM25 CLIP clusters across the protein coding genes reveals increased frequency in 5’ and 3’ UTRs. **(f,g)** RNA-immunoprecipitation (RIP) validates CLIP-seq data and PRY/SPRY as the RNA-interacting domain. HeLa cells were mock transfected, transfected with T7-TRIM25 or T7-TRIM25ΔRBD and the extracts immunoprecipitated with anti-T7 antibody. **(f)** qRT-PCR results from RIP assay for selected protein coding and non-coding RNAs. The values are relative to those obtained from mock transfected cells, which were set to 1. The mean and standard deviations (SD) of three independent biological replicates are shown. **(g)** qRT-PCR results from RIP assay for selected miRs. The values are relative to these obtained from mock transfected cells, which were set to 1. The mean and standard deviations of three independent biological replicates are shown.

To determine the TRIM25 binding sequence preference, we used the pyMotif program from the pyCRAC package and WebLogo on identified clusters to search for enriched k-mers. While no single strong consensus motif was identified, we found that TRIM25 prefers sequences rich in G and C (Fig. 4b). This was confirmed by a bioinformatics analysis showing a significantly higher GC content in TRIM25 clusters compared to random transcripts or random exons (p-values for both comparisons were 2.2e-16) (Fig. 4c). When we clustered our consensus motifs we found some discrete, G and C-rich short motifs (Fig. S6b). In the future, it will be interesting to check if any of these motifs contribute towards TRIM25’s RNA-binding specificity. Next, we assessed the distribution of TRIM25 binding in mRNAs. This revealed that most binding sites are in exons (56%), with the 3’UTR also prominently represented (23%) (Fig. 4d). Furthermore, we observed a noticeable enrichment of TRIM25 clusters just after the stop codon (Fig. 4e). TRIM25 binds to its own transcript, which could be biologically important (Figure S3b). Intriguingly, even though TRIM25 is localized to both nucleus and cytoplasm (Choudhury et al., 2014) intron binding events were not common in CLIP-seq analysis. This could be demonstrated on TRIM25 mRNA target – c-Myc. Both high throughput CLIP-seq and TRIM25 RNA Immunoprecipitation (RIP) followed by targeted amplification of c-Myc transcript showed that TRIM25 predominantly interacts with mature mRNA (Figure S3c-S3d). Nevertheless, TRIM25 interacts with many RNA-binding proteins (Fig. 3) and they could provide a platform for interactions with other targets at the level of pre-mRNAs.

To validate the CLIP-seq experiments, we performed RIP followed by qRT-PCR of selected TRIM25 targets in HeLa cells overexpressing either wild-type T7-TRIM25 or T7-TRIM25ΔRBD (Fig. 4f-4g). RIP qRT-PCR with the wild-type TRIM25 showed significant enrichment of tested mRNAs (Fig. 4f) and miRNAs (Fig. 4g). Intriguingly, the T7-TRIM25 RIP signal on *TRIM25* mRNA was much higher than that on other substrates (Fig. 4f). This could not be explained by increased levels of *TRIM25* mRNA, which was upregulated only three-fold when compared to the mock-transfected control (data not shown). Importantly, T7-TRIM25ΔRBD failed to precipitate any RNAs apart from *TRIM25* mRNA itself (Fig. 4f). These results could be explained by the binding of the T7 antibody to the nascent TRIM25, which was still associated with polyribosomes and *TRIM25* mRNA. As the T7 antibody could co-IP many nascent TRIM25 peptides, which were still associated with the *TRIM25* mRNA, the RIP signal would be increased. Altogether, these results uncover the first TRIM25/RNA interactome. This suggests that TRIM25 RNA-related functions may extend well beyond the regulation of pre-let-7 uridylation toward other aspects of RNA metabolism. Finally, these data clearly demonstrate that one of the main TRIM25 targets is its own mRNA.

To gain further insights into TRIM25’s role in RNA metabolism we have used qRT-PCR to analyze mRNA levels of selected targets (the same ones as assayed by RIP in Fig. 4f-g) and ZAP mRNA in the wild type and TRIM25 KO cells. The majority of analyzed transcripts did not display substantial difference between wild type and KO cells (Figure S7a-S7b). ZAP and RNU12 transcripts were moderately upregulated in the TRIM25 KO cells when compared with the wild type ones. To see if TRIM25 could regulate stability of transcripts we blocked pol II transcription by Actinomycin D and assayed the levels of selected RNAs by qRT-PCR in various time points (Figure S7c-S7e). No statistically significant difference between wild type and the TRIM25 KO cells was observed in the stability of ZAP and c-Myc transcripts (Figure S7c-e). This suggests that the steady-state ZAP mRNA level change seen in the TRIM25 KO cells is due to transcriptional control. This is not surprising as a recent study revealed that TRIM25 is a potent transcriptional regulator (Walsh et al., 2017). Further in-depth and high throughput experiments will be needed to fully understand the potential role of TRIM25 in RNA metabolism.

### Binding of TRIM25 to its mRNA augments TRIM25 ubiquitination

To uncover the RNA-related function of TRIM25 and the reason behind binding to its own mRNA, we focused on the well-documented molecular event of TRIM25 K117 autoubiquitination (Inn et al., 2011). We confirmed that TRIM25 undergoes autoubiquitination by co-overexpression and co-IP of T7-tagged TRIM25 together with HA-tagged ubiquitin (Figure S8a-S8b). This was also confirmed by the *in vitro* ubiquitination experiments described below. Next, we asked if binding of TRIM25 to its own mRNA augmented TRIM25 ubiquitination. To address this, we overexpressed EGFP-TRIM25 or EGFP-TRIM25ΔRBD in HeLa cells and assayed for TRIM25 ubiquitination. We used EGFP fusion proteins to distinguish overexpressed and endogenous TRIM25. The overexpression of EGFP-TRIM25 led to the appearance of modified EGFP-TRIM25 (Fig. 5a-5b). However, upon EGFP-TRIM25ΔRBD overexpression, the intensity of the modified band was noticeably reduced (Fig. 5a-5b). Of note, we had to use more EGFP-TRIM25ΔRBD plasmid to achieve protein expression comparable to that of EGFP-TRIM25. Surprisingly, overexpression of EGFP-TRIM25, but not EGFP-TRIM25ΔRBD, led to significant accumulation of modified forms of endogenous TRIM25, with much higher ratios of the modified forms compared to the non-modified form (Fig. 5a-5b). These results suggest that the intact RBD is required for efficient ubiquitination. They also show that endogenous TRIM25 can be modified by EGFP-TRIM25 to a much greater extent than EGFP-TRIM25 modifies itself. One important difference between endogenous *TRIM25* mRNA and the mRNA coding for EGFP-TRIM25 is the presence of 5’ and 3’ UTRs, which were shown to bind TRIM25 in the CLIP-seq experiments (Figure S3b).

**Fig. 5.**
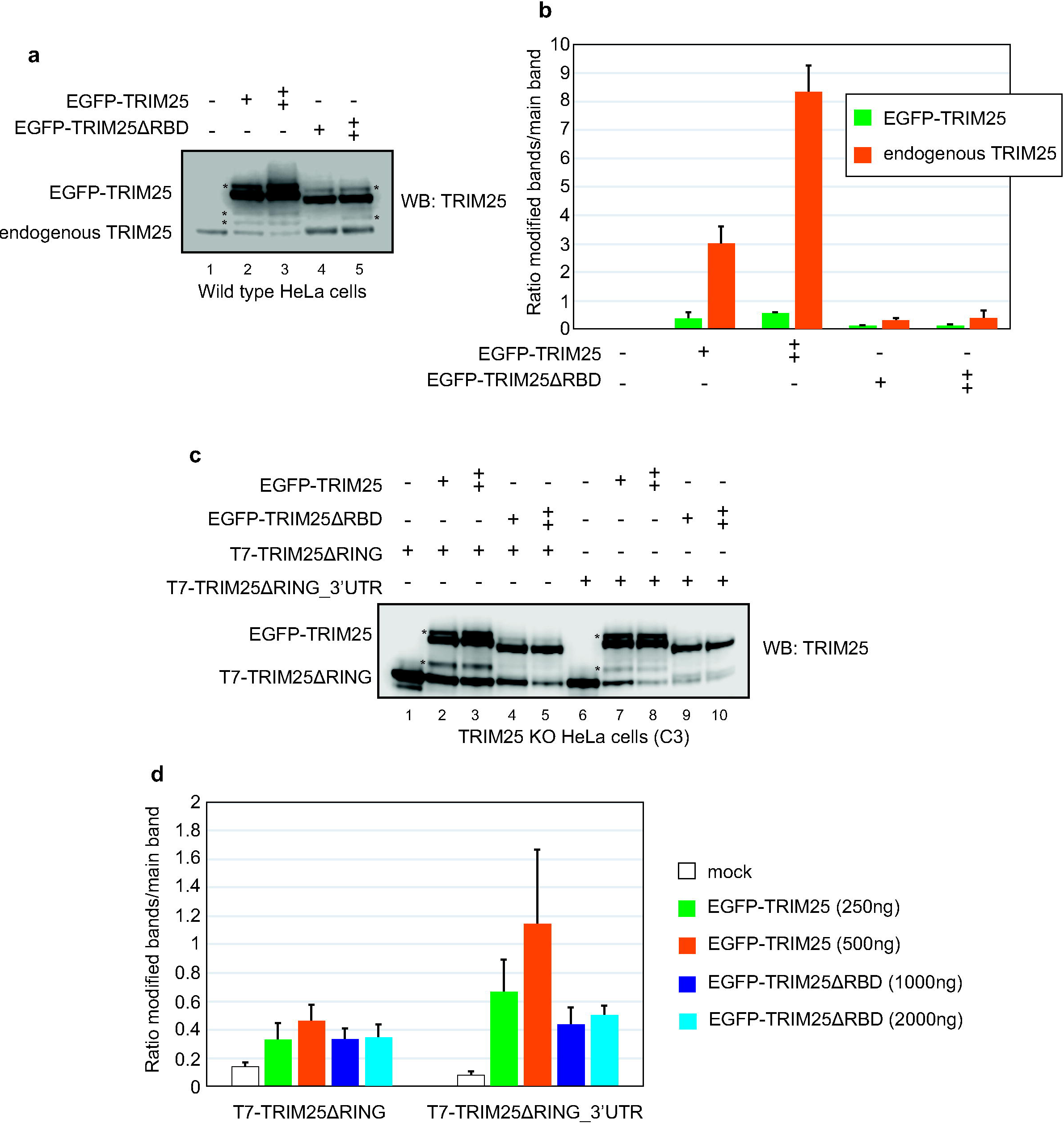
TRIM25 binding to the RNA augments TRIM25 ubiquitination. **(a)** Western blot analysis of HeLa cells transfected with EGFP-TRIM25 or EGFP-TRIM25ΔRBD. Lane 1 shows mock transfected control. Lanes 2 and 3 represent results of transfection with increasing amounts of plasmid coding for EGFP-TRIM25 (250 ng and 500 ng). Lanes 4 and 5 represent results of transfection with increasing amounts of plasmid coding for EGFP-TRIM25ΔRBD (1000 ng and 2000 ng). **(b)** Quantification of western blot results shown in **(a)**. The values show the ratios between the intensity of modified vs main bands. The results are derived from three independent biological replicates and the mean and standard deviations are shown. **(c)** Western blot analysis of TRIM25 KO HeLa cells transfected with plasmids coding for T7-TRIM25ΔRING (250 ng) or T7-TRIM25ΔRING_3’UTR (500 ng) as well as increasing amounts of plasmids coding for EGFP-TRIM25 or EGFP-TRIM25ΔRBD. Lane 1 shows T7-TRIM25ΔRING control. Lanes 2 and 3 represent results of transfections with T7-TRIM25ΔRING and increasing amounts of EGFP-TRIM25 (250ng and 500ng). Lanes 4 and 5 show results of transfection with T7-TRIM25ΔRING and increasing amounts of EGFP-TRIM25ΔRBD (1000 ng and 2000 ng). Lane 6 shows T7-TRIM25ΔRING_3’UTR control. Lanes 7 and 8 represent results of transfections with T7-TRIM25ΔRING_3’UTR and increasing amounts of EGFP-TRIM25 (250ng and 500ng). Lanes 9 and 10 show results of transfection with T7-TRIM25ΔRING_3’UTR and increasing amounts of EGFP-TRIM25ΔRBD (1000 ng and 2000 ng). **(d)** Quantification of western blot results shown in **(c)**. The values show the ratios between the intensity of modified vs main bands. The results are derived from three independent biological replicates and the mean and standard deviations are shown.

To determine if the *TRIM25* 3’UTR can contribute to ubiquitination of TRIM25, we used a catalytically dead T7-TRIM25ΔRING construct, which cannot undergo autoubiquitination, and transfected it into TRIM25 KO cells. We also used a T7-TRIM25ΔRING_3’UTR construct that contained the *TRIM25* 3’UTR. Notably, we had to use more T7-TRIM25ΔRING_3’UTR plasmid to achieve protein expression comparable to that of T7-TRIM25ΔRING (Fig. 5c). This is most likely connected to microRNA/3’UTR-mediated control of protein expression. As predicted, T7-TRIM25ΔRING does not support efficient autoubiquitination (Fig. 5c). However, upon EGFP-TRIM25 overexpression, we noticed the appearance of ubiquitinated forms of T7-TRIM25ΔRING (Fig. 5c-5d). We could not detect abundant ubiquitination bands in T7-TRIM25ΔRING when EGFP-TRIM25ΔRBD was co-overexpressed (Fig. 5c-5d). Even so, the ratio between the ubiquitinated and main bands still showed a positive value, most likely due to weaker expression of T7-TRIM25ΔRING in the presence of EGFP-TRIM25ΔRBD and to background noise. Importantly, when we co-overexpressed the T7-TRIM25ΔRING_3’UTR construct with EGFP-TRIM25, we observed a noticeable increase in the ratio of ubiquitinated vs non-ubiquitinated bands for TRIM25ΔRING, when compared with the T7-TRIM25ΔRING construct without the 3’UTR (Fig. 5c-5d). To prove that the observed modification of TRIM25 was specific to K117 we performed similar analyses with T7-TRIM25ΔRING_K117R constructs. We did not observe modification of T7-TRIM25ΔRING_K117R in the absence or presence of EGFP-TRIM25 (Figure S8c). Altogether, these results strongly suggest that RNA elements such as the 3’UTR, as well as TRIM25 binding to the RNA, contribute to TRIM25 ubiquitination.

### RNA is necessary for efficient TRIM25 ubiquitination *in vitro*

To test if RNA is necessary for efficient TRIM25 ligase activity, we performed *in vitro* ubiquitination assays with various TRIM25 constructs (Fig. 6a). Incubation of T7-TRIM25 or T7-TRIM25ΔRBD IPed from HeLa cells with recombinant proteins from the ubiquitin pathway (Ube2D3, UBE1, and Ubiquitin) revealed efficient, time-dependent ubiquitination of TRIM25 but not TRIM25ΔRBD (Fig. 6b). Residual ubiquitination of TRIM25ΔRBD could arise from the direct transfer of ubiquitin from Ube2D3 to the substrate. Next, to show that the observed ubiquitination was specific and dependent on the intrinsic activity of TRIM25, we assayed T7-TRIM25ΔRING and T7-TRIM25K117R. Neither T7-TRIM25ΔRING nor T7-TRIM25K117R was efficiently ubiquitinated compared to wild-type T7-TRIM25 (Fig. 6c). This confirmed the specificity of the *in vitro* ubiquitination assay. Finally, to see if TRIM25 ubiquitin-ligase activity was dependent on RNA, we performed the ubiquitination assay in the absence or presence of RNase A/T1. Remarkably, treatment with RNase A/T1 severely inhibited *in vitro* ubiquitination of T7-TRIM25 (Fig. 6d). Interestingly, RNase A/T1 treated reactions still supported Tim25 monoubiquitination. This could be due to the direct transfer of ubiquitin from Ube2D to TRIM25. These results strongly indicate that RNA constitutes an important component of the TRIM25 ubiquitin ligase activity.

**Fig. 6.**
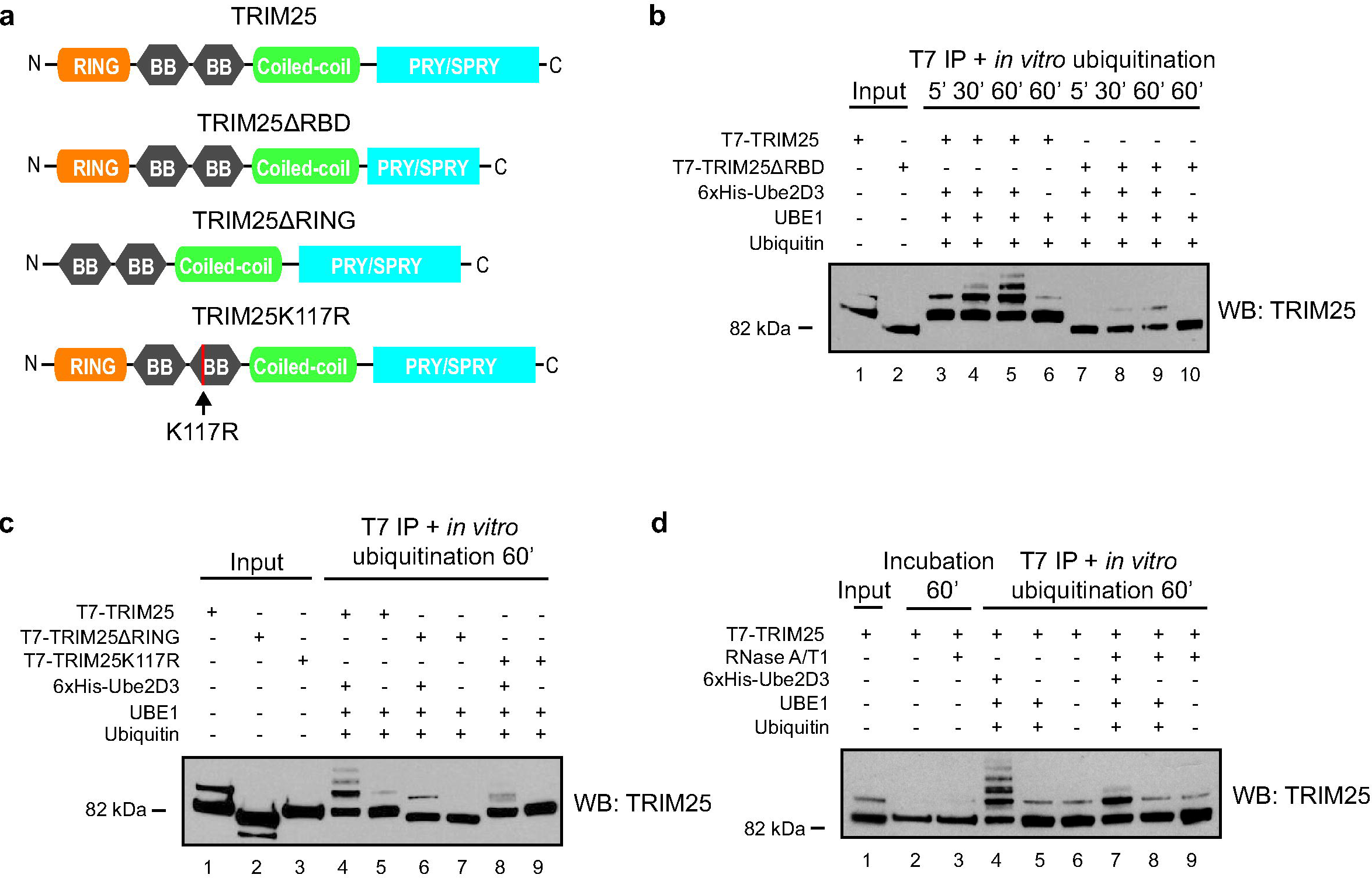
*In vitro* ubiquitination shows that RNA is important for TRIM25 E3 ubiquitin ligase activity. **(a)** Domain architecture of wild type TRIM25, TRIM25 deletions mutants and TRIM25 K117R mutant. **(b)** T7-TRIM25 but not T7-TRIM25ΔRDB is efficiently ubiquitinated *in vitro.* Ubiquitination assays were performed on IPed T7-TRIM25 or T7-TRIM25ΔRDB from HeLa cell extracts in the presence of recombinant purified 6xHis-Ube2D3, UBE1 and ubiquitin. Lanes 1 and 2 show IPed T7-tagged proteins. Lanes 3, 4 and 5 show incubation of T7-TRIM25 with 6xHis-Ube2D3, UBE1 and ubiquitin for 5, 30 and 60 minutes, respectively. Lane 6 represents 60 minutes’ reaction with T7-TRIM25, UBE1 and ubiquitin. Lanes 7, 8 and 9 show incubation of T7-TRIM25ΔRDB with 6xHis-Ube2D3, UBE1 and ubiquitin for 5, 30 and 60 minutes, respectively. Lane 10 represents 60 minutes’ reaction with T7-TRIM25ΔRDB, UBE1 and ubiquitin. The proteins were analyzed by western blotting with anti-TRIM25 antibody. **(c)** Neither T7-TRIM25ΔRING nor T7-TRIM25K117R are efficiently ubiquitinated *in vitro.* Ubiquitination assays preformed on IPed T7-TRIM25, T7-TRIM25ΔRING or T7-TRIM25K117R from HeLa cell extracts in the presence of recombinant purified 6xHis-Ube2D3, UBE1 and ubiquitin. Lanes 1, 2 and 3 show IPed T7-tagged proteins. Lanes 4, 6 and 8 represent 60 minutes’ reaction with T7-tagged IPed proteins, 6xHis-Ube2D3, UBE1 and ubiquitin. Lanes 5, 7 and 9 represent 60 minutes’ reaction with T7-tagged IPed proteins, UBE1 and ubiquitin. The proteins were analyzed by western blotting with anti-TRIM25 antibody. **(d)** RNase A/T1 treatment significantly reduces efficiency of T7-TRIM25 ubqiutinaiton. Lane 1 shows IPed T7-TRIM25. Lanes 2 and 3 represent control 60 minutes’ incubation of HeLa cell extracts in the absence and presence of RNases, respectively. Lanes 4 and 7 show reactions with IPed T7-TRIM25, 6xHis-Ube2D3, UBE1 and ubiquitin in the absence and presence of RNases, respectively. Lanes 5 and 8 show reactions with IPed T7-TRIM25, UBE1 and ubiquitin in the absence and presence of RNases, respectively. Lanes 6 and 9 show reactions with IPed T7-TRIM25 in the absence and presence of RNases, respectively. The proteins were analyzed by western blotting with anti-TRIM25 antibody.

### Homologous sequences from TRIM proteins rescue the TRIM25 RNA-binding and ubiquitination activities

To further verify our findings and test whether other TRIM proteins with PRY/SPRY domains could have RNA-binding potential, we prepared constructs encoding chimeric proteins, where the TRIM25 RBD was substituted with homologous sequences from several TRIM proteins (TRIM5, TRIM21, TRIM27 and TRIM65) (Fig. 7a). Moreover, we designed a TRIM25 construct with a scrambled amino acid sequence of the TRIM25 RBD (Fig. 7a). The chimeric TRIM25 protein constructs expressed in HeLa cells showed various levels of modification (Fig. 7b). Importantly, neither T7-TRIM25ΔRBD nor T7-TRIM25RBD_src was ubiquitinated (Fig. 7b). To see if these results translated to RNA-binding efficiency, we performed RNA pull-down assays with pre-let-7a-1 in HeLa cell extracts. Control assays confirmed that T7-TRIM25 could but T7-TRIM25ΔRBD could not bind RNA (Fig. 7c). All chimeric TRIM25 proteins, apart from T7-TRIM25ΔRBD_src, showed efficient binding to RNA (Fig. 7d-7e). Notably, we did not observe modified bands for TRIM25 in the RNA pull-down assays because the total protein amounts loaded on the gel were one order of magnitude lower than those in the overexpression experiments. To our surprise, there was a strong positive correlation (correlation coefficient 0.87, p-value 0.01) between protein modification and RNA pull-down efficiencies (Fig. 7f). Altogether, these results imply that RNA-binding is an important element in TRIM25 ubiquitination efficiency and suggest that many other TRIM proteins with the PRY/SPRY domain could harbor yet uncharacterized RNA-binding potential.

**Fig. 7.**
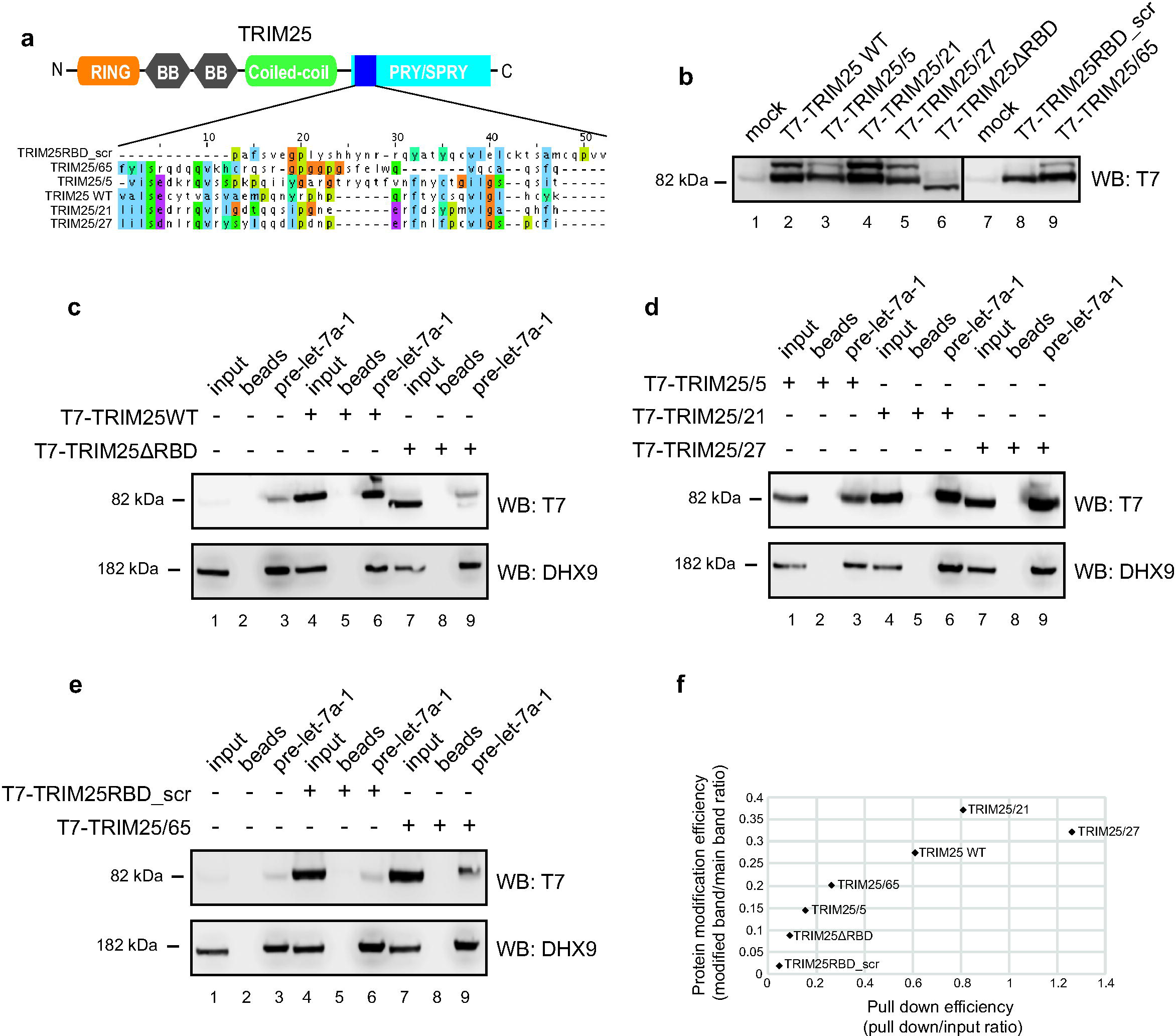
Chimeric TRIM25 proteins display correlated RNA-binding and ubiquitination activities. **(a)** Domain architecture of wild type TRIM25 with indicated RBD and chimeric homologous sequences introduced from other TRIM proteins as well as scrambled TRIM25 RBD sequence (aligned with Clustal Omega). **(b)** Western blot analysis of T7-TRIM25 chimeric proteins. Lanes 1 and 7 show blot of mock transfected HeLa cells. Lanes 2, 3, 4, 5, 6, 8 and 9 represent western blot analysis of overexpressed T7-TRIM25, T7-TRIM25/5, T7-TRIM25/21, T7-TRIM25/27, T7-TRIM25ΔRDB, T7-TRIM25RBD_scr and T7-TRIM25/65, respectively. **(c)** Western blot analyses against T7 and DHX9 of pre-let-7a-1 pull downs with HeLa cell extracts overexpressing T7-TRIM25 or T7-TRIM25ΔRBD. Lanes 1, 4 and 7 represent 4% (40 μg) of the loading controls (input). Lanes 2, 5 and 8 represent beads only pull downs. Lanes 3, 6 and 9 show pre-let-7a-1 pull down. Note that T7 antibody is highlighting an unspecific band visible in lanes 3 and 8. **(d)** Western blot analyses against T7 and DHX9 of pre-let-7a-1 pull downs with HeLa cell extracts overexpressing T7-TRIM25/5, T7-TRIM25/21 or T7-TRIM25/27. Lanes 1, 4 and 7 represent 4% (40 μg) of the loading controls (input). Lanes 2, 5 and 8 represent beads only pull downs. Lanes 3, 6 and 9 show pre-let-7a-1 pull down. **(e)** Western blot analyses against T7 and DHX9 of pre-let-7a-1 pull downs with HeLa cell extracts overexpressing T7-TRIM25RBD_scr or T7-TRIM25/65. Lanes 1, 4 and 7 represent 4% (40 μg) of the loading controls (input). Lanes 2, 5 and 8 represent beads only pull downs. Lanes 3, 6 and 9 show pre-let-7a-1 pull down. Note that T7 antibody is highlighting an unspecific band visible in lanes 3 and 6. **(f)** Graph displaying a positive person correlation coefficient (0.87) between TRIM25 ubiquitination efficiency (obtained by dividing signal intensities of modified protein bands by main protein bands **(b)**) and pull down efficiency (obtained by dividing signal intensities of pull down band to input **(c,d and e)**.

### RNA is necessary for efficient ubiquitination of ZAP by TRIM25 *in vitro*

To test if RNA was necessary for efficient ubiquitination of ZAP, a physiologically important TRIM25 target, we performed *in vitro* ubiquitination assays of ZAP (ZAPS) with the TRIM25 constructs described earlier (Fig. 6a). Incubation of T7-ZAP with T7-TRIM25 or T7-TRIM25ΔRBD IPed from HeLa cells with recombinant proteins from the ubiquitin pathway revealed efficient time-dependent ubiquitination of ZAP by TRIM25 but not TRIM25ΔRBD (Fig. 8b). Notably, to see efficient ubiquitination of ZAP we had to use higher concentrations of recombinant proteins from the ubiquitin pathway than those used for TRIM25 autoubiquitination experiments. As a result, TRIM25 autoubiquitination was much more pronounced in experiments with ZAP (Fig. 8a) than seen in the case of TRIM25 only assays (Fig. 6b). Furthermore, it has been recently reported that ZAP enhances TRIM25 autoubiquitination (Li et al., 2017). Next, to show that the observed ZAP ubiquitination was specific and dependent on the activity of TRIM25, we assayed T7-TRIM25ΔRING and T7-TRIM25K117R. TRIM25ΔRING but not T7-TRIM25K117R lost the ability to ubiquitinate T7-ZAP (Fig. 8b). Intriguingly, we observed autoubiquitination of T7-TRIM25K117R, which could be explained by increased concentrations of the proteins from the ubiquitination pathway and ZAP stimulatory activity towards TRIM25 (Li et al., 2017). Finally, to see if TRIM25 E3 ubiquitin ligase activity towards ZAP was dependent on RNA, we performed the ubiquitination assay in the absence or presence of RNase A/T1. Importantly, the treatment with RNase A/T1 severely inhibited *in vitro* T7-ZAP ubiquitination by T7-TRIM25 (Fig. 8c). These results strongly indicate that TRIM25 uses RNA as a scaffold for efficient ubiquitination of its protein targets.

**Fig. 8.**
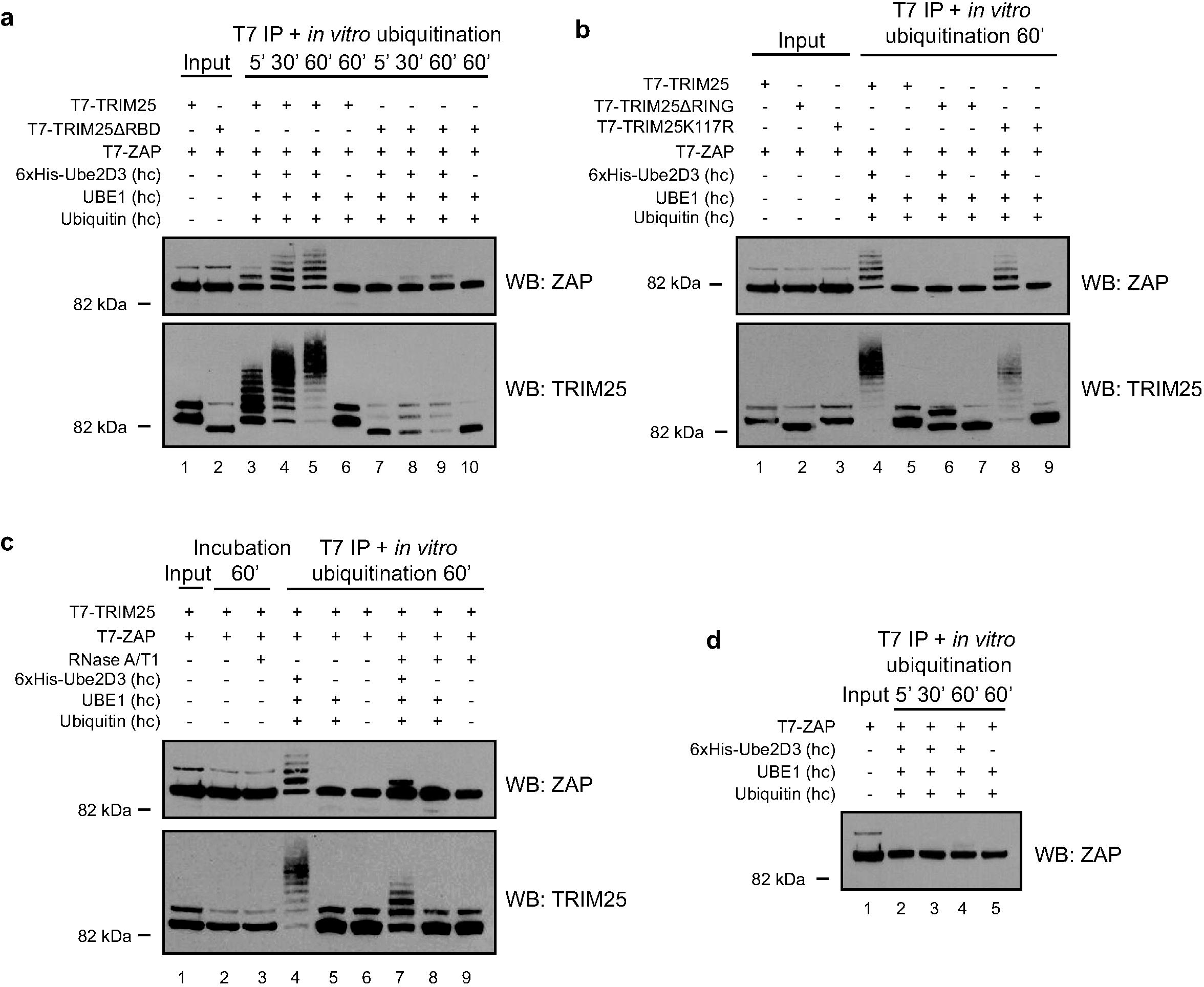
RNA is a scaffold for TRIM25-mediated ubiquitination of ZAP. **(a)** T7-TRIM25 but not T7-TRIM25ΔRDB efficiently ubiquitinated T7-ZAP protein *in vitro.* Ubiquitination assays were performed on IPed T7-ZAP with T7-TRIM25 or T7-TRIM25ΔRDB from HeLa cell extracts in the presence of recombinant purified 6xHis-Ube2D3, UBE1 and ubiquitin (the concentrations of proteins from the ubiquitin pathway was higher (hc) than in the experiments presented in Fig 6). Lanes 1 and 2 show IPed T7-tagged proteins. Lanes 3, 4 and 5 show incubation of T7-TRIM25 and T7-ZAP with 6xHis-Ube2D3, UBE1 and ubiquitin for 5, 30 and 60 minutes, respectively. Lane 6 represents 60 minutes’ reaction with T7-TRIM25, T7-ZAP, UBE1 and ubiquitin. Lanes 7, 8 and 9 show incubation of T7-TRIM25ΔRDB and T-7 ZAP with 6xHis-Ube2D3, UBE1 and ubiquitin for 5, 30 and 60 minutes, respectively. Lane 10 represents 60 minutes’ reaction with T7-TRIM25ΔRDB, T7-ZAP, UBE1 and ubiquitin. The proteins were analyzed by western blotting with anti-ZAP and anti-TRIM25 antibodies. **(b)** T7-TRIM25ΔRING but not T7-TRIM25K117R are defective in ZAP ubiquitination. Ubiquitination assays preformed on IPed T-ZAP with T7-TRIM25, T7-TRIM25ΔRING or T7-TRIM25K117R from HeLa cell extracts in the presence of recombinant purified 6xHis-Ube2D3, UBE1 and ubiquitin. Lanes 1, 2 and 3 show IPed T7-tagged proteins. Lanes 4, 6 and 8 represent 60 minutes’ reaction with T7-tagged IPed proteins, 6xHis-Ube2D3, UBE1 and ubiquitin. Lanes 5, 7 and 9 represent 60 minutes’ reaction with T7-tagged IPed proteins, UBE1 and ubiquitin. The proteins were analyzed by western blotting with anti-ZAP and anti-TRIM25 antibodies. **(c)** RNase A/T1 treatment substantially reduces efficiency of TRIM25-mediated ubqiutinaiton of ZAP. Lane 1 shows IPed T7-TRIM25 and T7-ZAP. Lanes 2 and 3 represent control 60 minutes’ incubation of HeLa cell extracts in the absence and presence of RNases, respectively. Lanes 4 and 7 show reactions with IPed T7-ZAP, T7-TRIM25, 6xHis-Ube2D3, UBE1 and ubiquitin in the absence and presence of RNases, respectively. Lanes 5 and 8 show reactions with IPed T7-ZAP, T7-TRIM25, UBE1 and ubiquitin in the absence and presence of RNases, respectively. Lanes 6 and 9 show reactions with IPed T7-ZAP and T7-TRIM25 in the absence and presence of RNases, respectively. The proteins were analyzed by western blotting with anti-ZAP and anti-TRIM25 antibodies. **(d)** Ubiquitination of T7-ZAP is mediated by T7-TRIM25. Control incubation of IPed T7-ZAP with 6xHis-Ube2D3, UBE1 and ubiquitin shows lack of ubiquitination. Lane 1 shows IPed T7-ZAP. Lanes 3, 4 and 5 show incubation of T7-ZAP with 6xHis-Ube2D3, UBE1 and ubiquitin for 5, 30 and 60 minutes, respectively. The proteins were analyzed by western blotting with anti-ZAP antibody.

## DISCUSSION

TRIM25 E3 ubiquitin ligase is a new RNA-binding protein with a role in Lin28a-mediated uridylation of pre-let-7 transcripts (Choudhury et al., 2014). Here, we revealed that the RNA-binding activity of TRIM25 is contained in its PRY/SPRY region. We also presented the first high-throughput analysis of the molecular interactomes of TRIM25. We showed that TRIM25 is a *bona fide* RNA-binding protein that associates with many proteins involved in RNA metabolism and interacts with numerous coding and non-coding transcripts. This suggests that TRIM25 could play a role in the regulation of RNA metabolism. Our data showed that TRIM25 binding to RNA and to the 3’UTR of *TRIM25* mRNA in particular can augment its autoubiquitination. Finally, we demonstrated that RNA is an important constituent of TRIM25’s ubiquitin ligase activity towards itself and its physiologically relevant protein target ZAP. Recently, ZAP was shown to suppress HIV-1 replication by binding CG-rich regions in viral RNAs (Takata et al., 2017). Intriguingly, the first paper that described ubiquitin conjugation to substrates showed that tRNA is required for selected ubiquitination reactions (Ferber and Ciechanover, 1986). We propose a model whereby the ubiquitin ligase activity of TRIM25 is substantially enhanced by using RNA as a scaffold (Fig. 9a). Alternatively, binding of TRIM25 to RNA could enhance its ubiquitin ligase activity by allosteric changes (Fig. 9b). These models should be verified by detailed structural analysis of the free and RNA-bound TRIM25. Finally, it is exciting to speculate that many other E3 ubiquitin ligases could depend on binding to RNA.

**Fig. 9.**
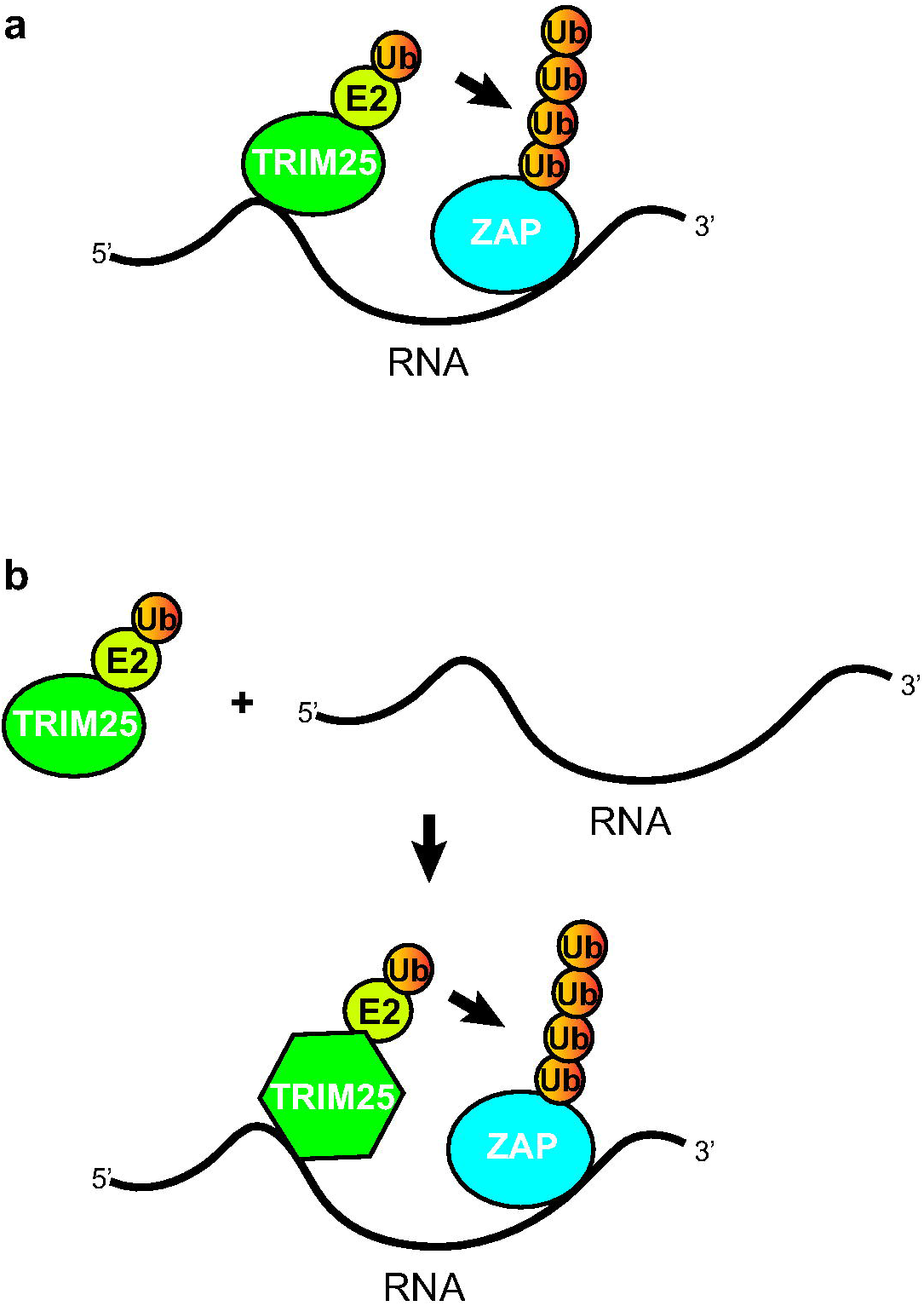
RNA is important for TRIM25’s E3 ubiquitin ligase function. **(a)** A model, whereby ubiquitin ligase activity of TRIM25 is substantially enhanced by using RNA as a scaffold **(b)** A model, whereby binding of TRIM25 to RNA could enhance its ubiquitin ligase activity by allosteric change.

Our results carry very important, functional implications given that the 3’UTRs are platforms for regulatory RNA-binding proteins and that the lengths of the 3' UTRs associated with specific mRNAs are variable. For example, cancer cells have mRNAs with much shorter 3' UTRs, which exhibit increased stability and produce more proteins in part through the loss of microRNA-mediated repression (Mayr and Bartel, 2009). Additionally, the shortening of 3' UTRs is associated with proliferating T cells but is not accompanied by a corresponding change in protein levels (Gruber et al., 2014). For these reasons, regulation of the lengths of the 3' UTRs could allow for tight control of the type and amount of TRIM25 that is associated with specific mRNAs and the subsequent ubiquitination/ISGylation of nascent peptides. This could be an important mechanism in the physiological and pathological control of protein stability and function. Importantly, TRIM25 also binds to the 5’UTRs of mRNAs and within coding regions. Thus, the RNA-related functions of TRIM25 might not be limited to 3’UTRs. This could result in different functions depending on where in the RNA the TRIM25 binds.

Previously, the CC domain of TRIM25 has been implicated in binding to RNA (Kwon et al., 2013). TRIM25 deletion mutants lacking the CC domain were unable to interact with RNA in the UV immunoprecipitation and T4 polynucleotide kinase assays (Kwon et al., 2013). In contrast, the mutant lacking the PRY/SPRY domain still retained RNA-binding ability. Surprisingly, the results from the proteome-wide analysis of RNA-binding domains in HeLa cells (Fig. 1b) (Castello et al., 2016), as well as the results obtained from the RNA pull-down (Fig.1e), EMSA (Fig. 1a,1c), co-IP (Fig. 3) and RIP (Fig. 4f-4g) assays, implicate the PRY/SPRY domain as the point of contact with the RNA. This discrepancy could arise from the fact that deletion of the CC domain from TRIM25 eliminates its dimerization (Li et al., 2014; Sanchez et al., 2014; Streich et al., 2013) and oligomerization (Koliopoulos et al., 2016) ability (Fig. 2c). While it is not univocally confirmed that TRIM25 needs to dimerize to bind RNA, such a phenomenon is widespread across many RNA-binding proteins (Feracci et al., 2016; Lunde et al., 2007). Additionally, the experiments presented in Kwon et al. were performed in wild-type mouse Embryonic Stem Cells (mESC), where levels of endogenous TRIM25 are particularly high (Kwon et al., 2013). This means that any ectopically overexpressed TRIM25 mutant (if it has the CC domain), including TRIM25ΔPRY/SPRY, will have a chance to heterodimerize with the wild-type, endogenous TRIM25. This could mask the requirements for a TRIM25 domain, such as PRY/SPRY, to bind to RNA. Our RNA pull-down data with the extracts from the TRIM25 KO cells clearly demonstrated that T7-TRIM25ΔPRY/SPRY could not bind RNA (Fig. 1g). Further structural characterization of the TRIM25/RNA complex should reveal novel features of this interaction and shed more light on this novel, putative RNA-binding domain.

Importantly, there are more than eighty TRIM proteins in the human genome (Versteeg et al., 2013), and they have been divided into 11 subfamilies based on their structural composition; forty-three of them have PRY/SPRY domains (subfamilies C-I and C-IV). Although there is a high degree of similarity between them, only TRIM25 (from TRIM proteins containing PRY/SPRY) was identified in HeLa (Castello et al., 2012) and mESC (Kwon et al., 2013) screens for proteins that bind to RNAs. This could reflect a lack of saturation of RNA-capture screens or lack of expression of many other PRY/SPRY-containing TRIM proteins in the cell lines used for these assays. Interestingly, another protein from the TRIM family, TRIM71, was shown to use its NHL, but not its CC, domain to bind RNA (Kwon et al., 2013; Loedige et al., 2013). Our results from the RNA pull-down experiments with chimeric TRIM25/TRIM5, TRIM25/TRIM21, TRIM25/TRIM27 and TRIM25/TRIM65 proteins indicate that other TRIM proteins with the PRY/SPRY domain could harbor yet undiscovered RNA-binding potential (Fig. 7). In this context TRIM5 was shown to generate K63 ubiquitin chains, which are required for restriction of retroviral reverse transcription (Fletcher et al., 2015). Thus, in the future, it will be important to assay other TRIM proteins to determine their RNA-binding activities. Notably, other E3 ubiquitin ligases, including Mex3b and Dizip3, have been shown to use RNA as a scaffold to mediate the ubiquitination of their substrates, which are bound to the same RNA (Yoon et al., 2013). Roquin is another example of an E3 ubiquitin ligase with a well-established RNA-binding ability (Leppek et al., 2013). Moreover, the BRACA1 E3 ubiquitin ligase, which is implicated in familial breast cancer, has been shown to interact with RNAs (Kawai and Amano, 2012). Altogether, these results strongly suggest the possibility of widespread roles of E3 ubiquitin ligases in the direct regulation of RNA metabolism.

TRIM25 is involved in normal development and disease (Heikel et al., 2016). For example, TRIM25 has been implicated in estrogen-dependent organ development (Inoue et al., 1993). Consequently, TRIM25 knockout mice present significantly underdeveloped uteri (Orimo et al., 1995; Orimo et al., 1999). Furthermore, TRIM25 plays a role in promoting proliferation of many cancer cell types, often in association with the estrogen response (Ikeda et al., 2000; Suzuki et al., 2005; Thomson et al., 2001; Urano et al., 2002). Recently, TRIM25 has been shown to be a master regulator of breast cancer metastasis (Walsh et al., 2017). Additionally, TRIM25 is known for its role in the Retinoic Acid-Inducible Gene 1 (RIG-I)-mediated interferon pathway (Ozato et al., 2008; Versteeg et al., 2013). Upon detection of viral RNAs with 5’ppp-RNA, RIG-I activates a signaling cascade that results in phosphorylation-mediated dimerization of the transcription factor IRF3, leading to type I interferon (interferon α and β) expression (Loo and Gale, 2011; Yoneyama et al., 2004). K63-linked ubiquitination of RIG-I by TRIM25 is required for an efficient interferon response (Gack et al., 2007). RIG-I can also be stimulated by unanchored polyubiquitin chains generated by TRIM25 (Zeng et al., 2010). This interaction is targeted by viruses to suppress the interferon response. For example, the influenza’s RNA-binding protein NS1 inhibits TRIM25-mediated ubiquitination of RIG-I by binding to the TRIM25 CC domain and disrupting its dimerization, allowing the virus to avoid the innate immune response (Gack et al., 2009). This inhibitory activity is significantly reduced in an NS1 mutant R38A/K41A, which cannot bind RNA (Donelan et al., 2003). This raises questions about a possible functional link between the RNA-binding activities of TRIM25 and NS1. Finally, SARS-CoV nucleocapsid protein (N-protein) has been shown to inhibit TRIM25 by binding to TRIM25’s PRY/SPRY domain (Hu et al., 2017). Considering our results, it is plausible that N-protein could directly inhibit the TRIM25 RNA binding activity. To fully understand the RNA-related role of TRIM25 in development and disease, it will be important to characterize TRIM25/RNA complexes, especially in the context of RIG-I activation by viral RNAs.

## CONCLUSIONS

In summary, we have presented the transcriptome and proteome-wide interactomes of a novel RNA-binding protein, TRIM25. We showed that the PRY/SPRY domain is responsible for the ability of TRIM25 to bind RNA. Finally, we have provided evidence that RNA is an important component in TRIM25’s ubiquitin ligase activity. A recent genome-wide bioinformatics screen of the human proteome identified more than 2600 proteins with RNA-binding capacity (Ghosh and Sowdhamini, 2016). Unravelling the RNA-related characteristics and functions of these proteins, including TRIM25, will undoubtedly reveal novel biologically and medically relevant molecular pathways.

## METHODS

### Electrophoretic mobility shift assay (EMSA)

EMSAs were performed with an end-labeled pre-let-7a-1 probe (50×10^3^ c.p.m. (counts per minute), ∼0.1 pmol) and the indicated amounts of proteins produced in E. coli. Binding reactions were performed in 16 μl reactions containing gel exclusion buffer (4.8 mM Tris pH 8, 144 mM NaCl, 0.96 mM DTT) supplemented with 3 mM MgCl_2_, 0.5 mM ATP, and 37.5 mM creatine phosphate. Reactions were incubated at 4°C for 1 h, followed by electrophoresis on a 6% (w/v) non-denaturing polyacrylamide gel. The signal was recorded with radiographic film or as exposed to a phosphoimaging screen and scanned on a FLA-5100 scanner (Fujifilm).

### RNA pull-down assay

RNA pull-down assays were performed as previously described (Choudhury et al., 2013), with slight modifications. In brief, total protein extracts were incubated with *in vitro* transcribed RNAs that were chemically coupled to beads (Michlewski and Caceres, 2010). The incubation was followed by a series of washes with buffer G (20 mM Tris [pH 7.5], 135 mM NaCl, 1.5 mM MgCl2, 10% (v/v) glycerol, 1 mM EDTA, 1 mM dithiothreitol, and 0.2 mM phenylmethanesulfonylfluoride). After the final wash, the proteins associated with the beads were analyzed by western blotting.

### Co-immunoprecipitation

Extracts prepared from HeLa cells transfected with plasmids expressing T7-tagged or EGFP-tagged proteins were incubated with protein A agarose with the anti-T7 antibody. The bound proteins were separated on a 4-12% SDS polyacrylamide gel and analyzed by western blotting. As indicated, some samples were treated with RNases A and T1 prior to loading on the gel. For mass spectrometry, the T7-bound beads were incubated with HeLa extracts grown in Light or Heavy R6K4 (13C-labeled arginine and 2D-labeled lysine) isotopes (Dundee Cell Products LM014/16). Next, the bound proteins were run on a 4-12% SDS polyacrylamide gel followed by in-gel digestion as described previously (Choudhury et al., 2013). Gene Ontology enrichment mapping was performed by functionally annotating the enriched proteins using DAVID (Huang da et al., 2009) and visualizing them with the Enrichment Map Plugin v2.1.0 (Merico et al., 2010) in Cytoscape (P-value cutoff = 0.001; FDR Q-value cutoff = 0.1; Jaccard coefficient = 0.25). Enriched protein sets are represented as nodes (red circles) connected by edges (green lines), with the sizes of nodes and edges representing the number of proteins in gene sets and the amount of overlap.

### CLIP-seq

The protocol was adapted from (Moore et al., 2014) using adapters and PCR oligonucleotides described in (Kilchert et al., 2015). In brief, two P100 dishes per sample were transfected with plasmid expressing T7-tagged TRIM25. Anti-T7 antibody was coupled to 50 μl Dynabeads per sample. Following immunoprecipitation, the RNA was dephosphorylated, the 3’ linker was ligated and radiolabeled and the 5’ linker was ligated on the beads. The 3’ linker reaction was performed in a total of 80 μl, with 1x PNK buffer, 1 μM 3’ adapter (3’ adapter in CLIP oligos), 800 U T4 RNA ligase 2 truncated K227Q, 80 U RNase out, and 10% PEG 8000. The 5’ linker reaction occurred in a total of 80 μl, with 1x PNK buffer, 1 mM ATP, 12.5 nM 5’ linker L5B (L5Ba-d in CLIP oligos), 80 U RNase OUT and 40 U T4 RNA ligase. The RNA was reverse transcribed (RT Oligo in CLIP oligos) and amplified by PCR (P5_Forward and P3_barcode Reverse BC1 or BC2) using LA Takara Taq. The libraries were analyzed on an Illumina Miseq system, and the data were processed using the pyCRAC software (Webb et al., 2014). TRIM25 binding motifs were analyzed using pyMotif and WebLogo on identified clusters. The distribution of TRIM25 CLIP clusters across the protein coding genes was performed using pyBinCollector. The correlation between the experiments was calculated by comparing the heights of the clusters in the three experiments. The correlation with total RNA abundance was performed by comparing the heights of the clusters common in all three experiments with the transcript abundance in the RNA-seq data derived from HeLa cells.

### RNA immunoprecipitation (RIP)

The RIP experiments were performed as in the CLIP-seq immunoprecipitations, except for cross-linking and RNase treatment. The bound RNAs were isolated by Trizol extraction, and RNAs were identified by qRT-PCR, using the SuperScript III Platinum SYBR Green One-Step qRT–PCR Kit (Life Technologies) following the manufacturer’s instructions on a Roche 480 LightCycler.

### *In vitro* ubiquitination

Immunoprecipitation was performed as described above. Following the final wash proteins bound to beads were incubated with Ube1, His-Ube2D3 and ubiquitin (UBPBio). 1 mg/ml Ube1, 2 mg/ml His-Ube2D3 and 50 mg/ml ubiquitin was used for TRIM25 autoubiquitination. 5 mg/ml Ube1, 10 mg/ml His-Ube2D3 and 250 mg/ml ubiquitin was used for ZAP ubiquitination. The reactions were carried out at 37C for 1 hour or as indicated and were stopped by addition of loading buffer and reducing agent. The protein ubiquitination was analysed by Western blot with anti-TRIM25 or anti-ZAP antibodies.

### Western blot

Total protein samples were resolved by standard NuPAGE SDS-PAGE electrophoresis with MOPS running buffer (Life Technologies) and transferred onto a nitrocellulose membrane. The membrane was blocked overnight at 4°C with 1:10 western blocking reagent (Roche) in Tris-buffered saline buffer with 0.1% Tween-20 (TBST). The following day, the membrane was incubated for 1 hr at room temperature with the primary antibody solution in 1:20 western blocking reagent diluted in TBST. Antibodies used included rabbit polyclonal anti-TRIM25 (1:1000, Abcam), rabbit polyclonal anti-DHX9 (1:1000, Protein-Tech), mouse monoclonal anti-tubulin (1:10000, Sigma-Aldrich), mouse monoclonal anti-T7 tag HRP conjugate (1:10000, Novagen), and mouse monoclonal anti-HA-probe HRP conjugate (1:1000, Santa Cruz). After washing in TBST, the blots were incubated with the appropriate secondary antibody conjugated to horseradish peroxidase and detected with the SuperSignal West Pico detection reagent (Thermo Scientific) or WesternSure ECL substrate (Licor). The membranes were stripped using the ReBlot Plus Strong Antibody Stripping Solution (Chemicon) equilibrated in water, blocked in 1:10 western blocking solution in TBST, and reprobed as described above.

### Cell culture, overexpression and Actinomycin D treatment

HeLa cells (American Type Culture Collection) were maintained in standard Dulbecco’s modified Eagle’s medium (DMEM) (Life Technologies), supplemented with 10% fetal bovine serum (Life Technologies). For SILAC mass spectrometry, cells were cultured in DMEM supplemented with Light or Heavy R6K4 (13C-labeled arginine and 2D-labeled lysine) isotopes (Dundee Cell Products LM014/16). Fragments encoding proteins of interest were cloned into the pCG T7 plasmid and transiently expressed in HeLa cells using Lipofectamine 2000 (Invitrogen). For RNA stability assays 10μg/mL Actinomycin D (Sigma) was added to cells and RNA extracted at the time points indicated using Tri Reagent (Sigma). Levels of RNAs of interest were determined by qRT-PCR using SuperScrip III Platinum SYBR Green One-Step qRT-PCR Kit (ThermoFisher). 200ng RNA was loaded per well for RNAs of interest and these levels were normalised to 18S Ribosomal RNA for which 1ng total RNA was loaded per well (1 in 200 dilution).

### Recombinant protein purification

The full-length human TRIM25 and RBD mutant were cloned into pET30a, which was transformed into BL21 gold cells. Bacteria was grown in Superbroth in the presence of kanamycin at 37°C until the OD_600_ reached 0.6. The bacteria were induced by 0.5 mM IPTG and incubated for 24 hours at 30°C with agitation. Pellets were lysed with a cell disruptor in lysis buffer (20 mM NaPPi pH 7.4, 500 mM NaCl, 5 mM MgCl2, protease inhibitor, DNase and RNase or benzonase), and lysates were loaded onto an IMAC HiTrap 1-ml FF column in binding buffer (20 mM NaPPi pH 7.4, 500 mM NaCl, 10 mM Imidazole) and eluted with increasing amounts of elution buffer (20 mM NaPPi pH 7.4, 500 mM NaCl, 500 mM Imidazole). Protein fractions were further purified with a gel exclusion column (Superdex 200 30/100 GL or HiLoad Superdex 200 16/60) in gel exclusion buffer (10 mM Tris pH 8.0, 300 mM NaCl, 2 mM DTT). Selected fractions were pooled and concentrated using Vivaspin 5 30000 MWCO (Sartorius Stedim UK Ltd.). His-Lin28s was produced and purified as specified earlier (Nowak et al., 2016).

### Thermal denaturation assay

Thermal denaturation assays were performed on a Bio-Rad IQ5 ICycler. Recombinant proteins (∼4 μM) were mixed with Sypro orange (x5) in a 50-μl reaction. Fluorescent readings were taken every 30 s between 20°C and 70°C. The T_m_ was determined from the maximum of the first derivative of the melting curve, and the mean T_m_ of three replicates was calculated for each protein.

### SEC-MALS

Mass measurements were performed on a Viscotek MALS-20 and VE3580 RI detector attached to an ÄKTA 10 Purifier with microflow components. Protein samples were run at approximately 0.75 mg/ml on a Superdex200 Increase 10/300 GL column, running at 1.0 ml/min in gel exclusion buffer (with 0.5 mM TCEP instead of DTT) with 100-μl injections.

### TRIM25 CRISPR/Cas9 knockout

HeLa cells were transfected with 200 ng GeneArt CRISPR Nuclease mRNA (Thermo Fisher) in addition to 50 ng of each of two sgRNAs (TRIM25 Left sgRNA – CCACGTTGCACAGCACCGTGTTC and TRIM25 Right sgRNA CTGCGGTCGCGCCTGGTAGACGG) targeting sequences in exon 1 of TRIM25. After 24 hours, cells were seeded to a 96-well plate such that there was on average 1 cell per well, and the cells were grown until colonies were visible by the naked eye. Next, cells were split into two 96-well plates, one of which was used for a dot blot analysis. For the dot blot analysis cells were washed twice in PBS before the addition of 30 μl Passive Lysis Buffer (Promega) per well and incubation for 15 minutes at room temperature. Two microliters of protein from each well was spotted directly onto a nitrocellulose membrane, followed by western blotting, as described above. Selected clones were seeded from the second 96-well plate into 6-well plates, grown and TRIM25 levels were validated by standard western blotting.

### CLIP oligos

L5Ba:

5'-invddT-ACACrGrArCrGrCrUrCrUrUrCrCrGrArUrCrUrNrNrNrArGrArGrCrN-OH-3’

L5Bb:

5'-invddT-ACACrGrArCrGrCrUrCrUrUrCrCrGrArUrCrUrNrNrNrGrUrGrArGrCrN-OH-3’

L5Bc:

5'-invddT-ACACrGrArCrGrCrUrCrUrUrCrCrGrArUrCrUrNrNrNrCrArCrUrArGrCrN-OH-3’

L5Bd:

5'-invddT-ACACrGrArCrGrCrUrCrUrUrCrCrGrArUrCrUrNrNrNrUrCrUrCrUrArGrCrN-OH-3'

3’ adapter:

5’App-NAGATCGGAAGAGCACACGTCTG-ddC 3’

RT Oligo:

5’-CAGACGTGTGCTCTTCCGATCT-3’

PCR Oligonucleotides:

P5_Forward:

5’-

AATGATACGGCGACCACCGAGATCTACACTCTTTCCCTACACGACGCTCTTCCGA

TCT-3’

P3_barcode_Reverse:

BC1:

5′CAAGCAGAAGACGGCATACGAGATCGTGATGTGACTGGAGTTCAGACGTGTGC TCTTCCGATCT-3′

(has indexing barcode IDX1)

BC2:

5′CAAGCAGAAGACGGCATACGAGATACATCGGTGACTGGAGTTCAGACGTGTGC TCTTCCGATCT-3′

(has indexing barcode IDX2)

## DECLARATIONS

### ACKNOWLEDGMENTS

We thank Hywel Dunn-Davies for providing a script for the analysis of the distribution of Clip-seq clusters in total RNA and in mRNA. We thank Finn Grey and Sam J. Wilson for the ZAP (ZC3HAV1) plasmid (Schoggins et al., 2011).

## AUTHORS’ CONTRIBUTIONS

GM conceived this study, GM, NRC, GH, MT and PK designed the experiments, NRC, GH, MT, PK and CS performed experiments, GM, NRC, GH, MT, PK, SW, SG, CS, JR and AC analyzed data, NRC and GM wrote the manuscript with input from other authors. All authors read and approved the final manuscript.

## ETHICS APPROVAL

Not applicable

## AVALIABILITY OF DATA AND MATERIALS

Not applicable

